# Dual effects of the small-conductance Ca^2+^-activated K^+^ current on human atrial electrophysiology and Ca^2+^-driven arrhythmogenesis: an *in silico* study

**DOI:** 10.1101/2023.06.16.545367

**Authors:** Nathaniel T. Herrera, Xianwei Zhang, Haibo Ni, Mary M. Maleckar, Jordi Heijman, Dobromir Dobrev, Eleonora Grandi, Stefano Morotti

**Affiliations:** Department of Pharmacology, University of California Davis, Davis, CA, USA; Computational Physiology Department, Simula Research Laboratory, Oslo, Norway; Department of Cardiology, Cardiovascular Research Institute Maastricht, Faculty of Health, Medicine, and Life Sciences, Maastricht University, Maastricht, The Netherlands; Institute of Pharmacology, West German Heart and Vascular Center, Faculty of Medicine, University Duisburg-Essen, Essen, Germany; Department of Medicine, Montreal Heart Institute and Université de Montréal, Montréal, QC, Canada; Department of Integrative Physiology, Baylor College of Medicine, Houston, TX, USA

**Author notes:** Correspondence to: Stefano Morotti, PhD, Department of Pharmacology, University of California Davis, Davis, CA, USA, Eleonora Grandi, PhD, Department of Pharmacology, University of California Davis, Davis, CA, USA.

**Keywords:** arrhythmia, atrial myocyte electrophysiology, atrial fibrillation, delayed afterdepolarizations, mathematical model, ion channels, reentry, spontaneous Ca^2+^release

## Abstract

By sensing changes in intracellular Ca^2+^, small-conductance Ca^2+^-activated K^+^ (SK) channels dynamically regulate the dynamics of the cardiac action potential (AP) on a beat-to-beat basis. Given their predominance in atria vs. ventricles, SK channels are considered a promising atrial-selective pharmacological target against atrial fibrillation (AF), the most common cardiac arrhythmia. However, the precise contribution of SK current (I_SK_) to atrial arrhythmogenesis is poorly understood, and may potentially involve different mechanisms that depend on species, heart rates, and degree of AF-induced atrial remodeling. Both reduced and enhanced I_SK_ have been linked to AF. Similarly, both SK channel up- and downregulation have been reported in chronic AF (cAF) vs. normal sinus rhythm (nSR) patient samples. Here, we use our multi-scale modeling framework to obtain mechanistic insights into the contribution of I_SK_ in human atrial myocyte electrophysiology. We simulate several protocols to quantify how I_SK_ modulation affects the regulation of AP duration (APD), Ca^2+^ transient, refractoriness, and occurrence of alternans and delayed afterdepolarizations (DADs). Our simulations show that I_SK_ activation shortens the APD and atrial effective refractory period, limits Ca^2+^ cycling, and slightly increases the propensity for alternans in both nSR and cAF conditions. We also show that increasing I_SK_ counteracts DAD development by reducing the coupling between transmembrane potential and intracellular Ca^2+^. Taken together, our results suggest that increasing I_SK_ in human atrial myocytes could promote reentry, while protecting against triggered activity. Depending on the leading arrhythmogenic mechanism, I_SK_ inhibition may thus be a beneficial or detrimental anti-AF strategy.

## Introduction

Atrial fibrillation (AF) is the most common clinical arrhythmia that significantly affects cardiovascular health worldwide (1, 2). It is characterized by rapid and irregular electrical activity and non-uniform electrical conduction leading to compromised atrial function. This impairment is linked to an enhanced risk of stroke and contributes to cardiovascular morbidity and mortality. The mechanisms underlying AF are complex, and involve both ectopic (triggered) activity from atrial foci (triggers) and a vulnerable substrate favoring impulse reentry through atrial tissue (3, 4). At the cellular level, focal ectopic/triggered activity is likely caused by early and delayed afterdepolarizations (EADs and DADs) or enhanced automaticity, while reentry is promoted by shortening of the action potential (AP) duration (APD), abbreviated atrial myocyte refractoriness (effective refractory period, ERP), and increased spatial and temporal heterogeneity (APD/ERP dispersion and alternans). In recent years, a large body of work has provided valuable insight in the molecular and cellular underpinnings of AF pathophysiology (5). Understanding the mechanisms underlying AF can facilitate targeted interventions, but at present pharmacological options demonstrate poor efficacy and safety concerns, and ablation is still the most effective clinical treatment (5). It is generally assumed that pharmacological treatment may be particularly suitable during the early stage of AF (i.e., paroxysmal AF), before significant electrical, structural, Ca^2+^ handling and contractile remodeling lead to long-standing persistent (chronic) AF (cAF) (6, 7). To avoid malignant adverse effects on ventricular electrophysiology, current research aims at exploiting atrio-ventricular electrophysiological differences to develop AF-selective strategies (8–11). For example, pharmacological approaches based on AF-selective blockade of the fast Na^+^ current (I_Na_) have been proposed (12, 13), as well as those targeting K^+^ channels primarily expressed in the atria (e.g., acetylcholine-sensitive and ultra-rapid K^+^ currents, I_K,ACh_ and I_Kur_) (9, 11).

Given their predominant expression in atria vs. ventricles, small-conductance Ca^2+^-activated K^+^ (SK) channels are emerging as a promising atrial-selective pharmacological target against AF (14–16). The family of SK channels consists of three members (17): SK1, SK2, and SK3, encoded by different genes (KCNN1, KCNN2, and KCNN3, respectively), and characterized by different sensitivity to the specific and potent channel pore blocker apamin (18). SK channels have been widely studied in the past two decades, since their identification in the heart and the assessment of their functional role in AP repolarization (17, 19). These initial studies showed that SK current (I_SK_) shortens atrial (but not ventricular) APD under physiological conditions. I_SK_ is modulated by Ca^2+^ (mediated by calmodulin tethered to the channel) (20), and can serve as an efficient process linking changes in intracellular Ca^2+^ cycling and sarcoplasmic reticulum (SR) Ca^2+^ release to transmembrane potential (E_m_) dynamics (21). Despite the lack of a voltage sensor in the channel, inwardly rectifying I_SK_-voltage relationships have been observed in many studies in both native and cloned channels from various species (22–25), and attributed to a voltage-dependent modulation of SK channel pore conductance by intracellular divalent (Ca^2+^ and Mg^2+^) cations (26, 27). Recent data revealed that SK channel trafficking also increases in a Ca^2+^-dependent manner (28, 29). These properties make SK channels of particular interest during fast pacing or tachy-arrhythmias (like AF), when Ca^2+^ accumulation is likely to enhance their activation. An increase in I_SK_ is expected to hasten repolarization, thus limiting APD and Ca^2+^ accumulation (a protective effect), but also to shortens the ERP, which may facilitate the formation of reentry circuits (proarrhythmic). Moreover, an altered dynamic APD regulation at fast rates may influence the induction of alternans (30), possibly promoting reentrant arrhythmias.

Indeed, both SK channels hyper-activity (24, 29, 31–38) and suppression (22, 25, 39) have been implicated in AF, likely through distinct mechanisms, and depending on species, heart rates, and disease etiology. Here, we aim to investigate the impact of modulating I_SK_ on the electrophysiology of human atrial myocytes and their vulnerability to Ca^2+^-driven arrhythmias using mathematical modeling and simulation, which have proven useful tool for gaining mechanistic insight into AF mechanisms, and developing novel antiarrhythmic pharmacological approaches (4, 40). Our findings contribute to a better quantitative understanding of the role of I_SK_ in various arrhythmia-provoking contexts, shed light on the underlying arrhythmia mechanisms, and provide insights into the potential proarrhythmic or protective effects of SK channel modulation in AF.

## Methods

### Human atrial cardiomyocyte simulations

To investigate the impact of SK channel modulation on atrial electrophysiology and arrhythmogenesis, we updated our established model of the human atrial myocyte in nSR and cAF conditions (41–43). We included into this cellular framework the I_SK_ formulation that we have recently developed based on recordings in human right-atrial cardiomyocytes from nSR and cAF patients (29). To simulate I_Na_, we used the formulation developed by Courtemanche *et al*. (44), and adjusted its maximal conductance to maintain the physiologic AP upstroke velocity. The formulation of the late Na^+^ current (I_NaL_) was modified by reducing the time constant of inactivation to reproduce experimental data at physiological temperature (45), as previously described (46). We also introduced a 21-mV rightward shift in the voltage-dependence of inactivation of I_NaL_ to match the half-activation potential of I_Na_ (44), and then adjusted I_NaL_ maximal conductance to reproduce the same current density during the AP. Lastly, we modified the cAF parameterization utilized in (43) by increasing the sensitivity of ryanodine receptor (RyR) for SR Ca^2+^ levels (i.e., 50% reduction in the EC_50_ for [Ca^2+^]_SR_-dependent activation of SR Ca^2+^ release) compared to the baseline nSR condition. Our default cAF model does not account for any AF-associated SK channel remodeling. However, we performed a separate set of simulations in which both SK channel conductance and Ca^2+^-sensitivity are altered as seen in AF (29, 47).

To gain a deeper understanding of the processes regulating Ca^2+^-driven arrhythmias, we also performed a set of simulations with our 3D model of the human atrial myocyte integrating membrane electrophysiology, spatially detailed Ca^2+^ handling, and variable tubular network (48, 49). We constructed ten different cardiomyocyte models generating different ultra-structures characterized by comparable overall low tubular densities, as previously described (48).

### Electrophysiologic protocols

Effect of modulation of the maximal conductance of SK channels (G_SK_) on AP and Ca^2+^ transient (CaT) properties at different pacing rates (i.e., between 0.5 and 4 Hz, with 0.5-Hz increments) were estimated at steady-state for both nSR and cAF. To investigate how G_SK_ changes affect atrial refractoriness, we determined the ERP simulating a S1-S2 premature stimulation protocol. First, for each condition studied, we identified the diastolic threshold of excitation (DTE). The S1 stimulus (5 ms in duration, 2-fold the DTE in amplitude) was applied at a basic cycle length (BCL) until steady-state. To determine the ERP, we then applied the premature S2 stimulus (same duration and amplitude as S1) at progressively smaller S1-S2 intervals by decrements of 5 ms. As previously described (50), the longest S1-S2 interval that failed to elicit an AP was taken as the local ERP (i.e., maximum upstroke velocity of ≥5 V/s and AP amplitude of ≥50% of that of the preceding AP elicited by S1).

We studied the role of G_SK_ modulation in Ca^2+^-driven atrial arrhythmias by investigating the determinants of alternans and DADs. The occurrence of alternans was assessed at steady state. By progressively increasing the stimulation frequency (1-ms decrements in BCL), we identified the maximum pacing thresholds required for the development of voltage (i.e., causing a ≥5 ms difference in APD estimated at E_m_ = -60 mV in subsequent beats) and Ca^2+^ (i.e., causing a ≥10 nM difference in CaT amplitude, CaT_amp_, in subsequent beats) driven alternans. Occurrence of DADs was assessed by stimulating the models at a fast rate for 10 s in the presence of 1 *μ*M Isoproterenol, and then pausing the stimulation. By progressively increasing the stimulation frequency (1-ms decrements in BCL), we identified the maximum pacing threshold required for the development of DADs (i.e., causing a voltage oscillation ≥10 mV during the pause).

We used the group of ten 3D models to assess the effect of G_SK_ modulation on spontaneous Ca^2+^ release events (SCRs) and consequent DADs at different pacing rates (i.e., 0.5, 1, 2, 3, 4, and 5 Hz). Each simulation consisted of a quiescent period of 4 s, a pacing period of 28 s to achieve steady-state, and a final quiescent period of 5 s for observation of oscillations in E_m_ and global [Ca^2+^]_i_. Whenever present, we quantified the amplitude of the voltage oscillation (ΔE_m_) by measuring the difference between the first peak and diastolic minimum during the final quiescent period. Then, we identified the corresponding SCR as the nearest [Ca^2+^]_i_ peak preceding the voltage oscillation, and determined its amplitude (ΔCa) by calculating the difference between the peak and diastolic minimum during the final quiescent period. We utilized cutoff values of 300 nM and 10 mV for oscillations in [Ca^2+^]_i_ and E_m_, respectively.

### Populations of human atrial cardiomyocyte models

Following an established approach (51), we built two populations of 1,000 atrial myocytes in nSR and cAF conditions by randomly perturbing the parameters of the Grandi *et al*. model listed in supplemental **Table S1** with log-normal scaling factors (σ = 0.1). Properties of AP and CaT, ERP, and maximum pacing thresholds for alternans and DADs were determined for each model variants in the two populations, as described above. To quantify the impact of parameter perturbations on the features analyzed, we performed multivariable linear and logistic regression analyses (51–53). Population size and parameter perturbation variance were chosen to ensure convergence of the results of these sensitivity analyses, as previously discussed (43, 54, 55).

### Numerical methods and code

Simulations of the updated Grandi *et al*. model were performed with a standard desktop using MATLAB (MathWorks, Natick, MA, USA), version R2022a. Simulations of the Zhang *et al*. 3D model (implemented in C++ and parallelized using OpenMP 5.1) were performed with a computing cluster with Intel Xeon CPU E5-2690 v4 at 2.60 GHz 28 CPUs (56 threads) + 132 GB; data analysis was performed with MATLAB using a standard desktop. Model codes are freely available on the authors’ website.

## Results

Our study utilized the Grandi *et al*. model of the human atrial cardiomyocyte in nSR and cAF (41), with subsequent modifications made as described in previous work (42, 43) and also detailed in the Methods section. We first quantified the impact of G_SK_ modulation (set to 50%, 100%, and 200% of the nominal value) on steady-state AP and CaT properties at varying stimulation frequencies (**Fig. 1**). In the absence of G_SK_ perturbation (i.e., 100% G_SK_), the cAF model displays shorter AP and ERP, and a more negative (polarized) resting E_m_ (RMP) compared to nSR. Despite the depressed cytosolic CaT, I_SK_ peak in cAF is comparable to that in nSR when the conductance does not undergo arrhythmia-driven remodeling (**Fig. 1A**). This is due to the fact that SK channel function is controlled by cleft and subsarcolemmal Ca^2+^ levels (i.e., not cytosolic), which rise both well above the K_d_ used to model Ca^2+^-activation of I_SK_ (K_d,SK_ = 350 nM) during the AP (not shown). The described differences between nSR and cAF AP and CaT features are maintained in the presence of G_SK_ perturbations. Our simulations show that increasing G_SK_ shortens the atrial AP and ERP, hyperpolarizes RMP, and reduces CaT_amp_ in both nSR and cAF myocytes (**Fig. 1B**). Simulations of the cAF model revealed the occurrence of alternans at fast pacing rates, independently of the G_SK_ value in use (**Fig. 1B**).

**Figure 1.**
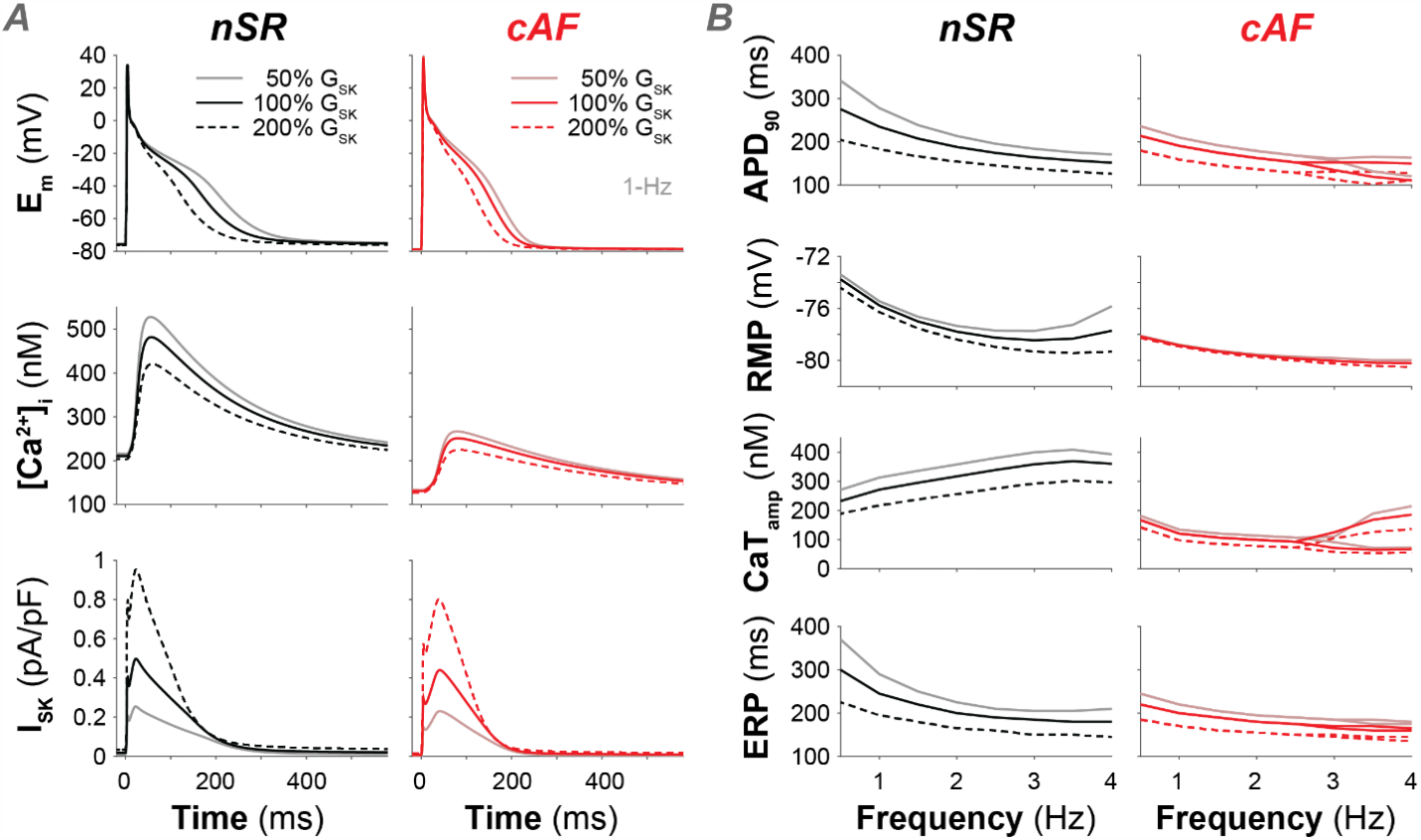
Increasing SK channel conductance shortens human atrial myocyte action potential duration (APD) and effective refractory period (ERP) in normal sinus rhythm (nSR) and chronic atrial fibrillation (cAF). A) Time course of membrane potential (E_m_), cytosolic Ca^2+^ concentration ([Ca^2+^]_i_), and SK current (I_SK_) in nSR and cAF myocytes during 1-Hz pacing for different values of SK channel maximal conductance (G_SK_). B) Effects of G_SK_ modulation on frequency-dependence of APD at 90% repolarization (APD_90_), resting membrane potential (RMP), Ca^2+^ transient amplitude (CaT_amp_), and ERP in nSR and cAF myocytes. Note that, when the cAF model develops alternans (i.e., at pacing rates of ≥3 Hz), values obtained at two subsequent alternating beats are reported.

To quantify the impact of small G_SK_ changes on human atrial myocyte electrophysiology, and assess the relative role of I_SK_ compared to other currents and transporters, we performed a parameter sensitivity analysis adopting an established methodology (51) based on the “population-of-models” approach (56). By randomly varying the values of maximal conductance and transport rate of ion channels and transporters (defined in **Table S1**), we generated two populations of atrial cells in nSR and cAF conditions reproducing the natural cell-to-cell variability in myocyte properties (see top panels in **Fig. 2**). We applied multivariable linear regression to quantify the sensitivity of APD, CaT_amp_, and ERP (assessed in each model variant at 1-Hz pacing) to changes in the perturbed model parameters (51). Our analysis revealed a predominant role of G_SK_ in modulating atrial myocyte APD and refractoriness (**Fig. 2A** and **C**), with a modest contribution to the regulation of CaT_amp_ (**Fig. 2B**). The coefficients associated to

**Figure 2.**
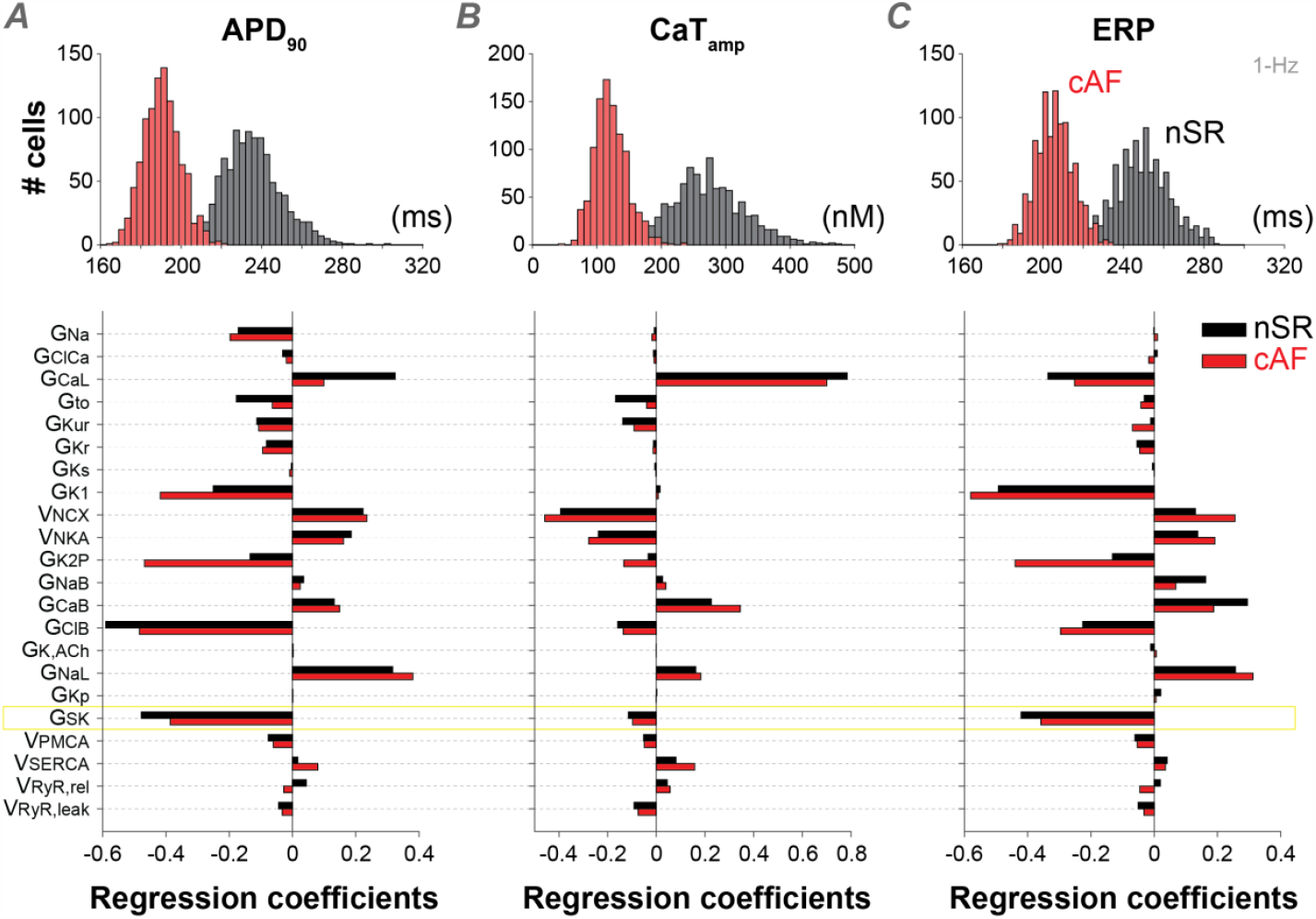
Sensitivity analysis reveals that SK channel conductance has a prominent role in modulating atrial myocyte action potential duration and effective refractory period. Top panels report histograms showing the distribution of APD_90_ (A), CaT_amp_ (B), and ERP (C) assessed at 1-Hz pacing in nSR and cAF populations of 1,000 models. Bottom panels report the results of linear regression analysis performed to quantify the sensitivity of APD_90_, CaT_amp_, and ERP (assessed at 1-Hz pacing) to changes in the listed individual parameters in the nSR and cAF models. Simulated data were obtained in nSR and cAF populations built upon baseline models with nominal SK channel maximal conductance (i.e., 100% G_SK_). Model variants exhibiting AP irregularities (1 in nSR, 4 in cAF) were excluded from linear regression analysis.

G_SK_ are negative, which indicates that increasing this parameter causes a reduction in APD, CaT_amp_, and ERP. The large amplitude of the G_SK_ coefficients for APD and ERP indicates that these features are highly sensitive to G_SK_ in both nSR and cAF, along with other K^+^ (i.e., basal inward rectifier and two-pore-domain currents, I_K1_ and I_K2P_) and background Cl^-^ currents. CaT_amp_ is less sensitive to K^+^ currents and primarily affected by changes in maximal conductance of L-type Ca^2+^ channels (G_CaL_) and transport rates of Na^+^/Ca^2+^ exchanger (v_NCX_) and Na^+^/K^+^ ATPase (v_NKA_), which are also positively correlated with APD changes, along with G_NaL_. Taken together, our results suggest that increased I_SK_ mainly causes APD and ERP shortening. These alterations have the potential to promote arrhythmogenesis as they increase the susceptibility to reentry, which can initiate arrhythmia within the atrial tissue and contribute to its persistence over time (4).

When pacing the cAF model at fast rates, our simulations revealed the development of alternans (**Fig. 1B**), another well-known contributor to reentrant arrhythmias (4). To identify the contribution of I_SK_ to this mechanism, we first assessed the maximal BCL at which alternans occur in the baseline nSR and cAF models (**Fig. 3A** and **B**). We separately determined BCL thresholds for development of beat-to-beat variations in APD (for simplicity called voltage-driven alternans) and CaT_amp_ (Ca^2+^-driven alternans). In agreement with the results shown in **Fig. 1**, BCL thresholds are longer in cAF vs. nSR (343 and 348 ms vs. 194 and 195 ms at 100% G_SK_ for voltage- and Ca^2+^-driven alternans, respectively), indicating that propensity for alternans is enhanced in disease. While G_SK_ changes minimally alter BCL thresholds in nSR (197/197 and 203/204 ms at 50% and 200% G_SK_, respectively, for voltage/Ca^2+^-driven alternans), increasing G_SK_ moderately elevates the BCL thresholds in cAF (330/334 and 352/358 ms at 50% and 200% G_SK_, respectively, for voltage/Ca^2+^-driven alternans). We then determined the BCL thresholds for voltage-driven alternans in each model variant of our nSR and cAF populations, and performed multivariable linear regression (51) to quantify how changes in model parameters alter this threshold (**Fig. 3C**). The positive regression coefficients determined for G_SK_ indicate that increasing this parameter prolongs the BCL for alternans in both nSR and cAF, increasing the vulnerability to reentry. However, the small amplitude of these coefficients (with respect to those of other coefficients) indicates that the contribution of I_SK_ to development of alternans is limited compared to that of other channels or transporters. Our results show that the parameters that are more influential for the pacing threshold for alternans in both nSR and cAF are G_CaL_ and the maximal rate of SR Ca^2+^ release through RyRs (v_RyR,rel_). Increasing these parameters would reduce the pacing threshold, thereby protecting against alternans.

**Figure 3.**
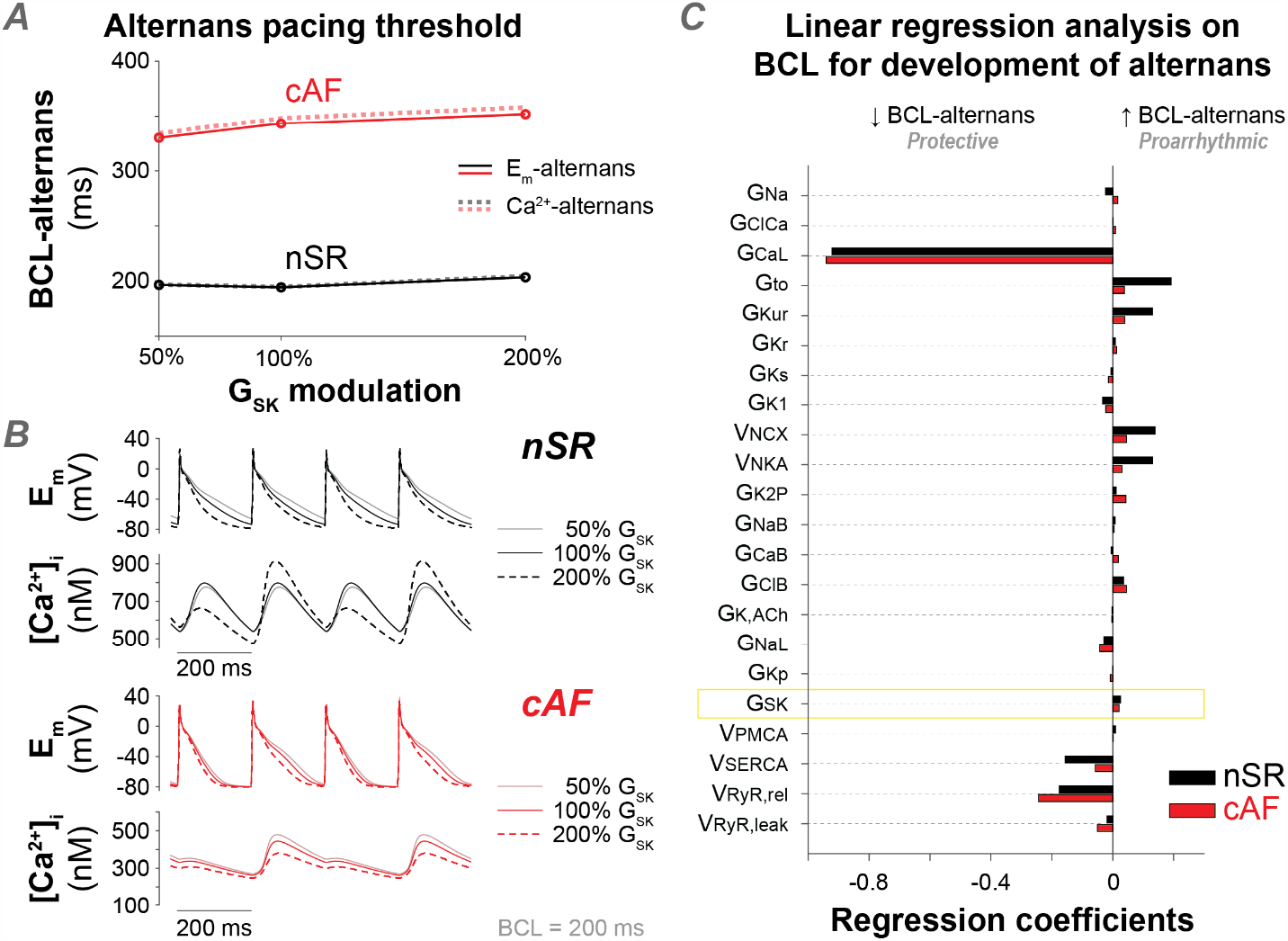
Increasing SK channel conductance modestly favors development of alternans in nSR and cAF myocytes. A) Effect of G_SK_ modulation on the maximal basic cycle length (BCL) inducing voltage-(solid lines) and Ca^2+^-driven (dotted lines) alternans in the baseline nSR and cAF models. B) Time course of membrane potential and cytosolic Ca^2+^ concentration obtained at steady-state in the baseline nSR and cAF models during 5-Hz pacing with variable SK channel maximal conductance. C) Results of linear regression analysis performed to quantify the sensitivity of maximal BCL for voltage-driven alternans development (BCL-alternans) to changes in the listed model parameters in nSR and cAF. Simulated data were obtained in populations of 1,000 nSR and cAF models built upon baseline models with nominal SK channel maximal conductance (i.e., 100% G_SK_). Model variants exhibiting alternans at 1-Hz pacing rate (1 in nSR, 4 in cAF) and model variants that never exhibited alternans (14 in nSR) were excluded from the linear regression analysis, which was performed analyzing the remaining 985 nSR and 996 cAF models.

We next assessed the involvement of I_SK_ in the development of DADs, a Ca^2+^-driven phenomenon that can cause triggered activity, thereby contributing to AF initiation (4). We first assessed the impact of G_SK_ modulation on the maximal BCL at which DADs occur in our baseline nSR and cAF models (**Fig. 4A** and **B**). BCL thresholds are higher in cAF vs. nSR (574 vs. 345 ms at G_SK_ 100%), confirming the disease-associated increase in DAD propensity.

**Figure 4.**
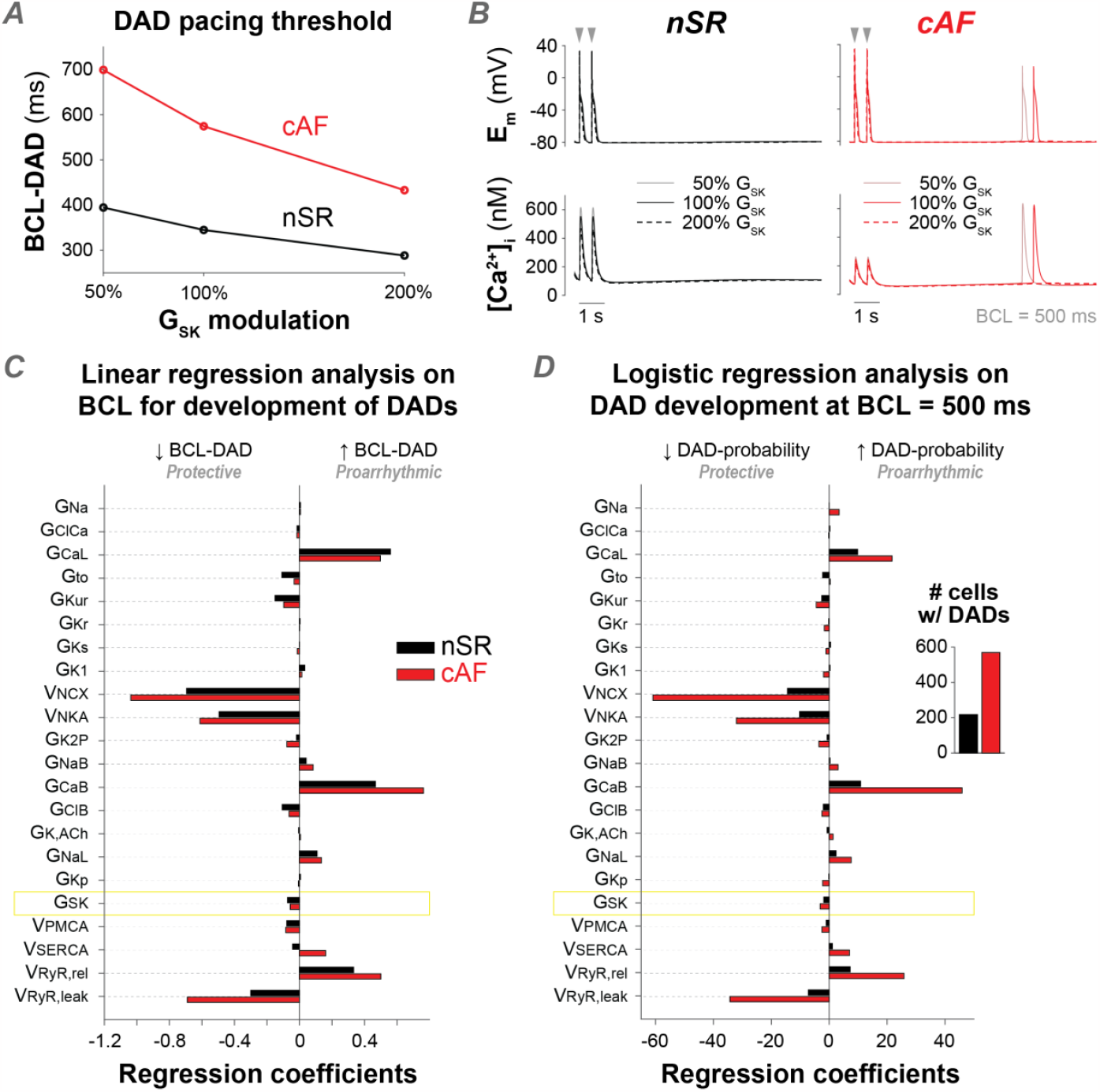
Increasing SK channel conductance protects against delayed afterdepolarization (DAD) development in nSR and cAF myocytes. A) Effect of G_SK_ modulation on the maximal basic cycle length (BCL) inducing DADs in the baseline nSR and cAF models. B) Time course of membrane potential and cytosolic Ca^2+^ concentration elicited in the baseline nSR and cAF models after pausing the electrical stimulation after 10 s of 2-Hz pacing in the presence of Isoproterenol (1 *μ*M) with variable G_SK_. Only the last two paced beats are reported in the figure. C) Results of linear regression analysis quantifying the sensitivity of maximal BCL for DAD development (BCL-DAD) to changes in the listed model parameters in nSR and cAF. D) Results of logistic regression analysis quantifying the impact of the listed model parameters on the probability of DAD development in nSR and cAF at 2-Hz pacing (i.e., BCL = 500 ms). Intercept terms b_0_ (determining the probability of DAD development in the absence of parameter perturbations) are -19.1 for nSR and +12.7 for cAF. The inset shows the number of model variants showing DADs during 2-Hz pacing within the nSR and cAF populations. Simulated data analyzed in C and D were obtained in populations of 1,000 nSR and cAF models built upon baseline models with nominal SK channel maximal conductance (i.e., 100% G_SK_). Model variants exhibiting DADs at 1-Hz pacing rate (129 in nSR, 422 in cAF) and model variants that never exhibited DADs (24 in nSR, 62 in cAF) were excluded from the linear regression analysis shown in C, which was performed analyzing the remaining 847 nSR and 516 cAF models.

Furthermore, increasing G_SK_ reduces the BCL thresholds in both nSR (394 and 288 ms at 50% and 200% G_SK_, respectively) and cAF (699 and 432 ms at 50% and 200% G_SK_, respectively), thereby protecting against arrhythmias. As described for alternans, we determined the BCL thresholds for DADs in each model in our populations, and performed linear regression (51) to quantify how changes in model parameters alter the propensity for DADs (**Fig. 4C**). The negative regression coefficients determined for G_SK_ indicate the protective effect of I_SK_ activation against DADs in both nSR and cAF (i.e., increasing G_SK_ shortens the BCL for DADs). Our analysis also highlights that the sensitivity to changes in G_SK_ is limited if compared to changes in other parameters that are directly involved in the regulation of Ca^2+^ cycling, such as G_CaL_, v_NCX_, v_NKA_, and v_RyR,rel_. We next separated our nSR and cAF populations in two sub-groups of model variants based on the development of DADs at 2-Hz pacing (i.e., models with a BCL threshold ≥500 ms). Using this cutoff, we identified 218 and 570 model variants exhibiting DADs within the nSR and cAF populations, respectively (**Fig. 4D**, inset). As previously described (53), we performed multivariable logistic regression to quantify how changes in model parameters affect the probability of DAD development in nSR and cAF. Our results show a negative correlation between G_SK_ modulation and DAD probability (**Fig. 4D**), confirming that I_SK_ activation is protective against DADs. This analysis also shows that modulation of G_SK_ has a smaller effect on DAD probability compared to changes in G_CaL_, v_NCX_, v_NKA_, and v_RyR,rel_, corroborating the findings obtained with linear regression (**Fig. 4C**).

To get mechanistic insight into the sub-cellular processes involved in the development of triggered activity, we simulated our recently-developed 3D model of the human atrial myocyte with spatially detailed Ca^2+^ cycling (48, 49). We constructed ten different cardiomyocyte models characterized by similar low density but different organization of the transverse tubular network, and assessed the relationship between the occurrence of SCRs and DADs and the pacing rate. In agreement with our previous results, we found that an increase in G_SK_ is protective against DADs (i.e., lower BCL threshold, see left panel in **Fig. 5A**). Interestingly, the BCL threshold for SCR is not affected by G_SK_ modulation (**Fig. 5A**, right), suggesting reduced coupling between E_m_ and [Ca^2+^]_i_ with increased G_SK_. To confirm this hypothesis, we analyzed both supra- and sub-threshold E_m_ and [Ca^2+^]_i_ oscillations obtained after stimulating the cells at a pacing frequency of 3 Hz (**Fig. 5B** and **C**). We found that increasing G_SK_ significantly reduces the amplitude of E_m_ oscillations (ΔE_m_). The amplitude of the [Ca^2+^]_i_ oscillations (ΔCa) is not different between 50% and 100% G_SK_, but is significantly reduced with 200% G_SK_. To assess changes in E_m_-Ca^2+^ coupling, we calculated the ΔE_m_/ΔCa ratio, and identified a significant reduction with an increase in G_SK_. Notably, we obtained similar results when investigating different pacing rates between 2 and 5 Hz (**Fig. S1**). Taken together, these results indicate that the protective effect of G_SK_ activation against development of DADs is due to a reduction of E_m_-Ca^2+^ coupling.

**Figure 5.**
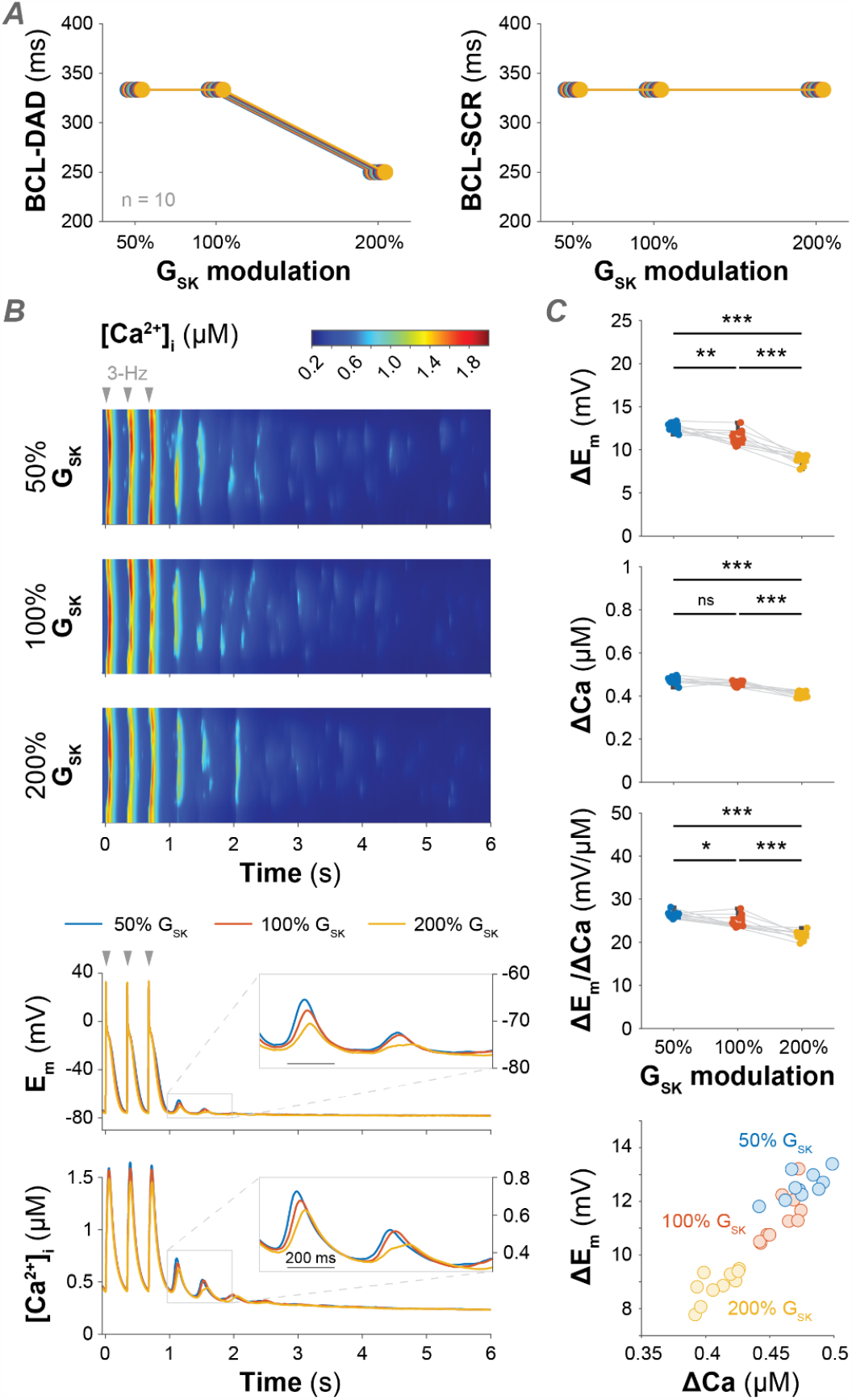
Increasing SK channel conductance protects against DADs by reducing the coupling between transmembrane potential and intracellular Ca^**2+**^. A) Effect of changes in G_SK_ on the maximal basic cycle length (BCL) inducing DADs and spontaneous Ca^2+^ release events (SCRs) in a human atrial myocyte model with three-dimensional Ca^2+^ diffusion. Thresholds for DADs and SCRs were set to 10 mV and 300 nM, respectively. The simulations were repeated for ten randomly generated tubular structures. B) Top panels show representative transverse line scans of local cytosolic Ca^2+^ concentration obtained pausing the electrical stimulation after a train of impulses at 3-Hz pacing for different G_SK_ values. The last three paced beats are reported in the figure. Bottom panels report the time course of corresponding global cytosolic Ca^2+^ concentration and membrane potential. C) Summary data resulting from simulating the protocol described in panel B in the 10 different tubular structures.Top panels show amplitude of E_m_ and [Ca^2+^]_i_ oscillations (ΔE_m_ and ΔCa), and the ΔE_m_/ΔCa ratio in function of different G_SK_ values. Statistical analysis was performed by one-way ANOVA with Bonferroni correction (***: p < 0.001; **: p < 0.01; *: p < 0.05; ns: not significant). ΔE_m_ and ΔCa are also compared against each other in the scatter plot reported in the bottom panel.

## Discussion

We performed a computational analysis aimed at unraveling the role of SK channels in human atrial myocyte electrophysiology and arrhythmogenesis. Using our established multi-scale modeling framework, we simulated various electrophysiology protocols to determine the impact of I_SK_ modulation on the regulation of APD, CaT, and ERP, as well as the occurrence of arrhythmogenic alternans and DADs. Our findings indicate that increased G_SK_ promotes APD and ERP shortening, while slightly increasing the propensity for alternans, thereby primarily promoting the development of an arrhythmogenic reentrant substrate. Our simulations also demonstrated that enhanced I_SK_ limits Ca^2+^ cycling and counteracts the occurrence of DADs, thus exerting a protective effect against triggered activity. This protective effect is due to a reduction in E_m_-[Ca^2+^]_i_ coupling when G_SK_ is increased. Overall, these results (summarized in **Fig. 6**) highlight the potential dual effect of targeting SK channels for the treatment of AF depending on the leading arrhythmogenic mechanism. These observations align with existing literature reporting that both reduced and enhanced I_SK_ may predispose to AF in animal models (22, 24, 25, 31–39). As discussed below, these contrasting outcomes of I_SK_ modulation can be attributed to species differences, variability in proarrhythmic protocols, as well as the extent and type of AF-induced atrial remodeling.

**Figure 6.**
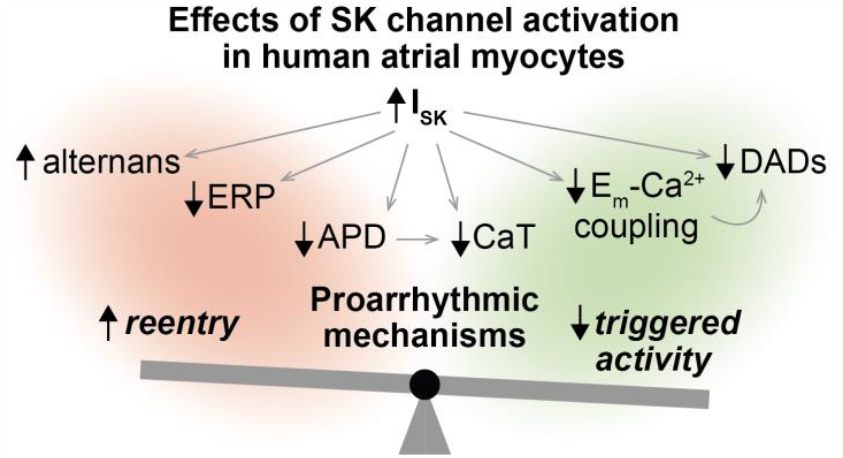
Dual effects of SK channel activation in human atrial myocytes. Our simulations indicate that increased SK current promotes *a)* APD and ERP shortening, *b)* slight increase in propensity for alternans, *c)* depressed Ca^2+^ cycling, *d)* decrease in coupling between transmembrane potential and intracellular Ca^2+^, and *e)* reduced propensity for DADs. While effects *a* and *b* promote the development of an arrhythmogenic reentrant substrate, effects *c, d*, and *e* counteract triggered activity.

### SK contribution to AF remodeling and arrhythmogenesis

Progression of AF is associated with extensive remodeling of ion channels and Ca^2+^ handling proteins (5). A genome-wide association study suggested a possible link between human lone AF and an intronic single nucleotide polymorphism in the KCNN3 gene (57, 58), but the precise role of SK channels in atrial arrhythmogenesis is still poorly defined (14–16). Indeed, both I_SK_ upregulation and inhibition have been associated with increased AF propensity. Ozgen *et al*. first showed that fast pacing increases SK2 channel expression in rabbit atria (59). Subsequent studies have confirmed this finding in other species (rat, guinea pig, dog, goat, horse), and showed the protective effect of pharmacological I_SK_ inhibition (24, 31, 32, 34–38), which can also be mediated by secondary effects on I_Na_ properties (60, 61). On the other hand, Hsueh *et al*. showed that I_SK_ block in dog prolongs APD, facilitates development of alternans, and is proarrhythmic (39). Reduction in SK2 and SK3 channel expression has been reported in a diabetic mouse model, resulting in APD prolongation and arrhythmias (25). Finally, increased

AF susceptibility was found in both SK2 knock-out mice (22) and mice over-expressing SK3 channels (33), the latter genotype being more prone to sudden cardiac death (62). The degree of AF-induced remodeling has also been shown to impact I_SK_ function, whereby studies in dog and human have shown that SK channel expression is reduced in cAF (23, 63–65), and I_SK_ block does not affect the APD. Other studies have instead reported increased I_SK_ in cAF (29, 47, 66). In our recent investigation, we did not find changes in SK channel expression at mRNA or protein level, but we demonstrated that I_SK_ is significantly increased in human cAF vs. nSR, due to enhanced Ca^2+^-dependent SK channel gating and membrane trafficking and targeting (29).

Given the complex regulation of I_SK_ in human data, we did not include any AF-associated change in SK channel function during cAF simulations. Adopting the same nominal G_SK_ used for nSR, we determined that I_SK_ plays a primary role in the regulation of APD and ERP in the human atrial myocyte in cAF (**Fig. 2**). When repeating the same analysis accounting for a 2-fold increase in G_SK_ in cAF vs. nSR (**Fig. S2**), as suggested by our functional experiments (29), we found, as expected, an increased relative sensitivity to G_SK_ changes (see cAF coefficients in **Fig. S2** vs. **Fig. 2**). Interestingly, under this assumption, I_SK_ becomes the more influential K^+^ current for regulation of atrial myocyte repolarization and refractoriness (surpassing I_K1_). In our cAF simulations, we also assumed unchanged Ca^2+^-dependent activation of I_SK_ compared to nSR. Experiments in atrial cardiomyocytes from patients have shown increased affinity for intracellular Ca^2+^ in cAF vs. nSR (47), which could contribute to I_SK_ upregulation in disease (29). Increasing Ca^2+^ affinity minimally impacts the results produced with the Grandi *et al*. model, especially in terms of regulation of APD, ERP, and CaT (**Fig. S3A**). As discussed earlier, this is because SK channel function is controlled by the levels of Ca^2+^ in cleft and subsarcolemmal compartments that, during the AP, rise well above the K_d_ values used here. Our data revealed minor changes in the propensity for alternans and DADs, with the former being facilitated and the latter being counteracted by decreased K_d,SK_ (**Fig. S3B** and **C**). The limited impact of increasing Ca^2+^ affinity is confirmed with the Zhang *et al*. 3D model (**Fig. S4**). While BCL threshold for SCRs are not affected, decreasing K_d,SK_ lowers the BCL threshold for DAD in two (out of ten) atrial myocyte models at nominal G_SK_ (see panels **A** in **Fig. S4** vs. **Fig. 5**). Taken together, these results suggest that enhanced G_SK_ and increased Ca^2+^ affinity can both contribute to SK channel gain of function and have a protective effect against cellular triggered activity. Based on these observations and preliminary data indicating increased I_SK_ in patients with paroxysmal AF (67), we speculate that I_SK_ upregulation can be an adaptive change to the increased propensity to DAD-mediated mediated cellular triggered activity that occurs during the early stage of AF, when focal firing is thought to contribute to the periodic re-initiation of AF in the absence of clinically-relevant ERP/APD abbreviation (68). As AF progresses toward a persistent (chronic) stage, SK channel gain of function turns into a maladaptive change that contributes to AF maintenance by causing reentry-promoting APD/ERP abbreviation (29, 69).

### Targeting SK channels in cAF patients

Despite the contrasting results reported in literature, inhibition of I_SK_ has emerged as a promising anti-AF strategy and is currently being pursued in a clinical trial (ID: NCT04571385) (70). Based on our findings, a successful completion of this trial would support the notion that the main mechanisms underlying AF is linked to reentry, rather than focal activity. However, the outcome of pharmacological treatment can be strongly affected by patients’ variability linked to stages of AF progression (discussed before), presence of comorbidities, and sex differences. For example, AF often coexists with heart failure (HF), leading to worse prognosis and more complicated patient management (71, 72). It has been shown that I_SK_ is upregulated in ventricular myocytes during HF (64, 65), where apamin can prolong the AP, but it is still not clear whether I_SK_ activation would either facilitate or counteract ventricular arrhythmias (73). Similar to our observations in atria, Terentyev *et al*. showed that upregulation of SK channels in HF attenuates DADs driven by spontaneous Ca^2+^ waves, thereby reducing triggered activity in ventricles (21). Bonilla *et al*. suggested that pharmacological I_SK_ inhibition in HF causes excessive AP prolongation, leading to repolarization instability and development of EADs (64). On the other hand, the modeling study by Kennedy *et al*. showed that I_SK_ activation produces arrhythmogenic alternans and instabilities in single cell and tissue simulations in ventricles (74). These observations suggest that SK channel inhibition in AF patients with concomitant HF might induce additional side effects at the ventricular level.

It is well known that sex differences in cardiac electrophysiology exist and can affect pathophysiology and response to treatment (75). This also applies to AF, where several differences have been identified in prevalence, clinical presentation, associated comorbidities, and therapy outcomes in females vs. males (76). Notably, the former are more likely to be diagnosed with AF at an earlier age, and the latter are more likely to have worse AF symptoms that typically last longer and are more severe (77). The mechanistic bases of these sex differences are still poorly understood, mostly because studies investigating sex-specific mechanisms of AF pathophysiology have so far been very limited. Experiments in rabbit ventricular myocytes revealed increased I_SK_ in female vs. male animals (78). If these sex differences were confirmed in human atria, SK channel modulation might induce larger relative changes in female vs. male myocytes, and if confirmed in human ventricles, SK inhibition could cause harmful AP/QT prolongation in females. Interestingly, no females (forty-seven healthy male volunteers) were recruited in the successful phase I clinical trial NCT04571385 designed to test safety and tolerability of AP30663 (70), an SK2 channel inhibitor that is showing antiarrhythmic efficacy (79).

### Limitations and possible extensions

In the present study, we have not explicitly considered the contribution of protein kinase A (PKA) and Ca^2+^/calmodulin-dependent protein kinase II (CaMKII) signaling. These pathways are important regulators of atrial myocyte function (80), and can synergistically promote atrial arrhythmogenesis (81). Since both PKA and CaMKII can affect SK channel function (29, 47), further studies are needed to explore the effects of I_SK_ modulation in the context of altered signaling in disease. We previously showed that the effects of I_SK_ inhibition on lengthening atrial myocyte AP and ERP are consistent between single cells and tissues (43). Moreover, our tissue simulations revealed that reduced I_K1_ promotes source-sink mismatch and triggered activity (81). We speculate that I_SK_ inhibition may lead to a similar outcome. Lastly, our atrial myocyte models were built assuming homogeneous distribution of SK channels through the plasmalemma (48). Experiments in rabbit ventricular myocytes suggested instead that SK channels are predominantly located near L-type Ca^2+^ and RyR2 channels (82). Further studies assessing how this spatial proximity influences local and global Ca^2+^ signaling and E_m_ dynamics are clearly warranted.

### Conclusions and future directions

Our study shows a dual effect of SK channel modulation in human atrial cardiomyocytes of both nSR and cAF. By counteracting reentry arrhythmias, I_SK_ inhibition may be suitable for treatment of cAF patients. This strategy, however, may cause an increase in vulnerability for cellular DADs, thus potentially promoting atrial ectopy. These observations are similar to those made for APD-prolonging class III agents. The use of these drugs that target the hERG K^+^ channel (responsible for the delayed rectifier K^+^ current I_Kr_) is constrained by the potential risk of inducing life-threatening ventricular arrhythmias through EADs (83). To better characterize benefits and risks of I_SK_ modulation in female vs. male patients, future research should investigate the contribution of SK channels to sex differences in atrial electrophysiology, remodeling, and arrhythmogenesis. In addition, efforts should be made to address the potential additional complications due to concomitant occurrence of HF, whereby pharmacological SK channel inhibition might induce adverse side effects in ventricles, potentially in part mediated by altered mitochondrial function (84). Integrative computational modeling frameworks allow comprehensive characterization of the effects of I_SK_ modulation (alone or in combination with other targets, as shown in (43, 85)) on male and female atrial and ventricular electrophysiology, thereby supporting the development of safe and effective therapeutical treatments against AF.

## Acknowledgements

The authors wish to thank Dr. Jussi Koivumäki for helpful discussion at the initial stages of this work.

## Authors Contributions

EG and SM conceived and designed the research; NTH, XZ, and HN performed *in silico* experiments and data analysis; NTH, XZ, HN, MMM, JH, DD, EG, and SM interpreted results of experiments, NTH and XZ prepared the figures; NTH and SM drafted the manuscript; XZ, HN, MMM, JH, DD, and EG edited and revised the manuscript; all authors approved the final version of manuscript.

