## SUPPLEMENTARY MATERIALS for "Dual effects of the small-conductance Ca^2+^-activated K^+^ current on human atrial electrophysiology and Ca^2+^-driven arrhythmogenesis: an *in silico* study"

[@] Correspondence to:

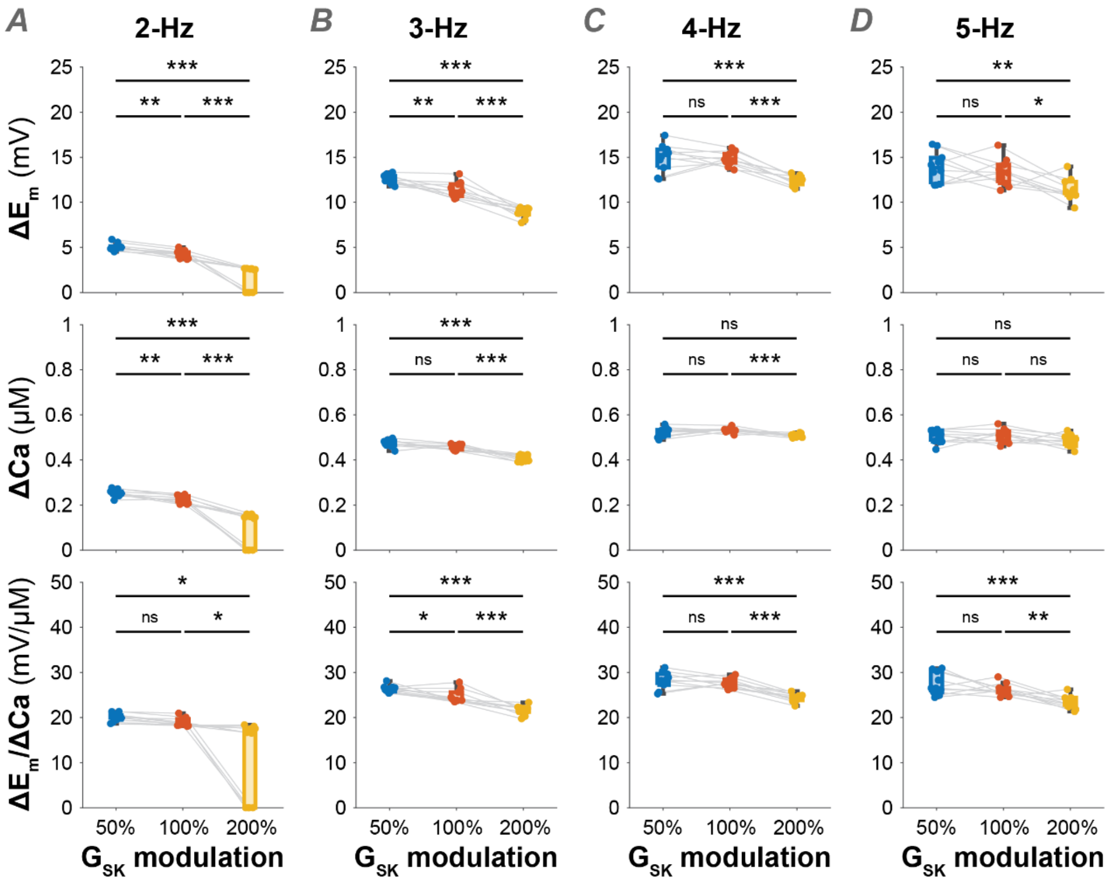


**Figure S1. Increasing SK channel conductance reduces the coupling between transmembrane potential and intracellular $Ca^{2+}$ over a broad range of pacing rates.** The analysis shown in **Fig. 5B** and **C** was performed varying the stimulation frequency. No voltage or $[Ca^{2+}]_i$ oscillations were observed at 0.5 and 1-Hz pacing rates. We report here the summary data describing amplitude of membrane potential ($E_m$) and cytosolic $Ca^{2+}$ concentration ($[Ca^{2+}]_i$) oscillations ($\Delta E_m$ and $\Delta$Ca), and the $\Delta E_m/\Delta$Ca ratio in function of different values of SK channel maximal conductance ($G_{SK}$) at 2 (1), 3 (B), 4 (C), and 5 Hz (D). Null values of $\Delta E_m$, $\Delta$Ca, and $\Delta E_m/\Delta$Ca shown for 200% $G_{SK}$ at 2-Hz pacing indicate absence of voltage or $[Ca^{2+}]_i$ oscillations. Statistical analysis was performed by one-way ANOVA with Bonferroni correction (***: $p < 0.001$; **: $p < 0.01$; *: $p < 0.05$; ns: not significant).

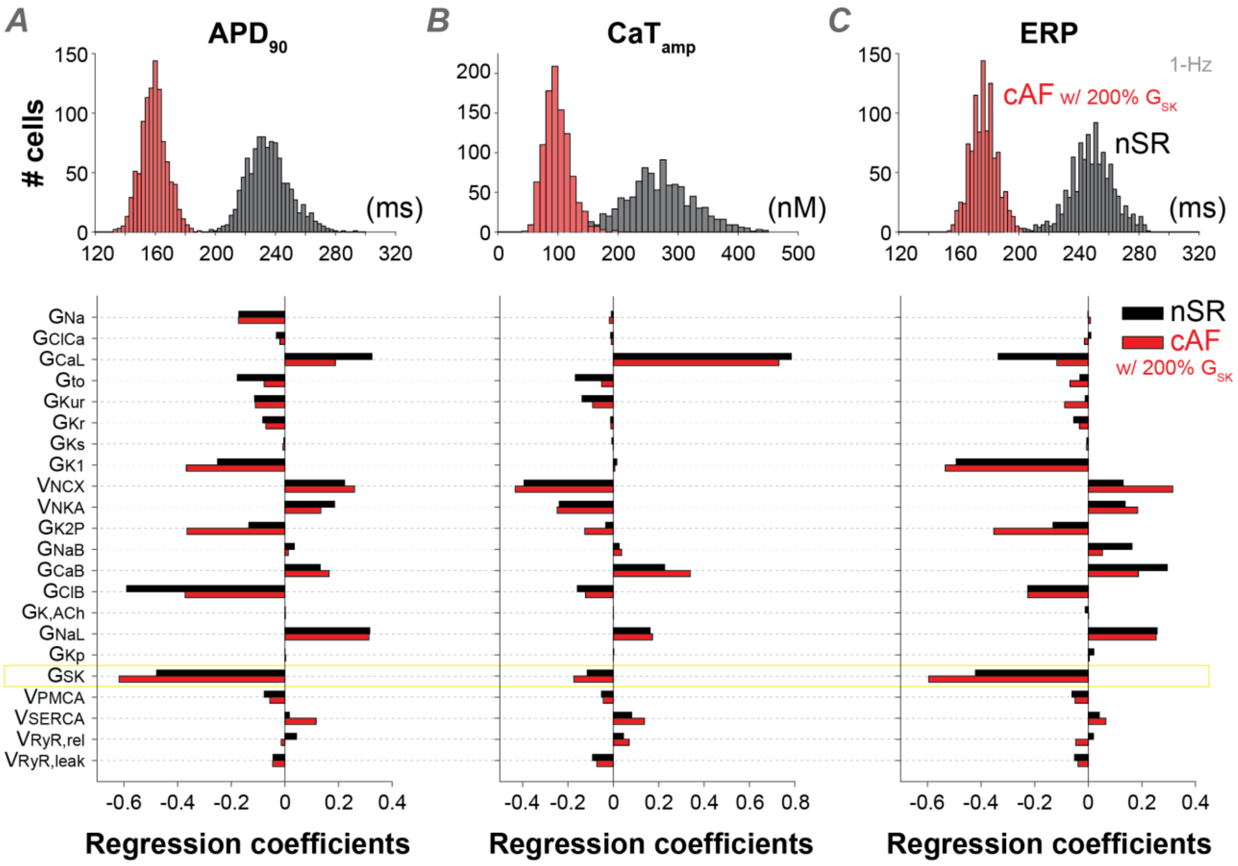


**Figure S2. Sensitivity analysis performed simulating increased SK expression in chronic atrial fibrillation (cAF) vs. normal sinus rhythm (nSR).** The analysis shown in **Fig. 2** was replicated assuming increased $G_{SK}$ in cAF vs. nSR myocytes. Top panels report histograms showing the distribution of action potential duration (APD) at 90% repolarization ($APD_{90}$, A), $Ca^{2+}$ transient amplitude ($CaT_{amp}$, B), and effective refractory period (ERP, C) assessed at 1-Hz pacing in nSR and cAF populations of 1,000 model variants. Bottom panels report the results of linear regression analysis performed to quantify the sensitivity of $APD_{90}$, $CaT_{amp}$, and ERP (assessed at 1-Hz pacing) to changes in the listed parameters in the nSR and cAF models. Simulated nSR data were obtained in a population of models built upon a baseline model with nominal SK channel maximal conductance (i.e., 100% $G_{SK}$). Simulated cAF data were obtained in a population of models built upon a baseline model with increased SK channel maximal conductance (i.e., 200% $G_{SK}$). Model variants exhibiting AP irregularities (1 in nSR, 4 in cAF) were excluded from the linear regression analysis.

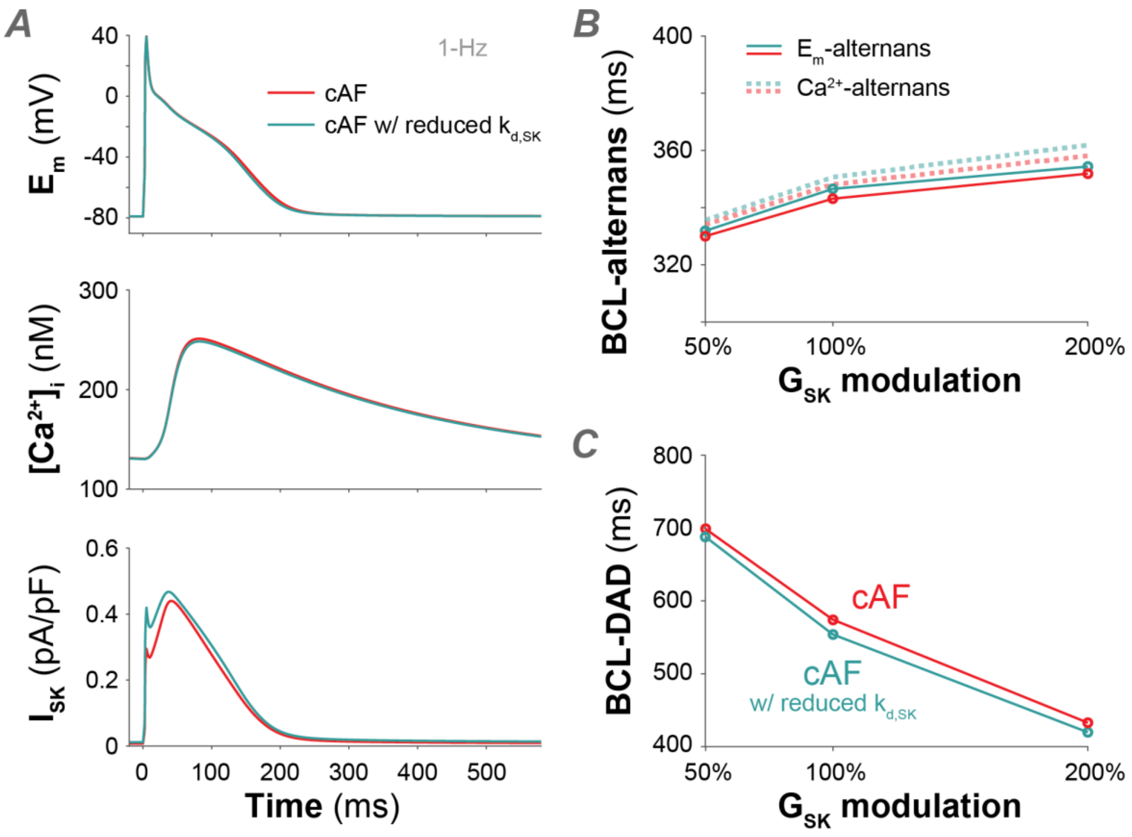


**Figure S3. Increased SK channel $Ca^{2+}$ affinity moderately affects atrial myocyte electrophysiology and arrhythmogenesis in cAF.** A) Time course of $E_m$, $[Ca^{2+}]_i$, and SK current ($I_{SK}$) elicited in cAF myocytes during 1-Hz pacing with nominal (i.e., $K_{d,SK}$ = 350 nM) or increased (i.e., $K_{d,SK}$ = 230 nM) SK channel affinity for intracellular $Ca^{2+}$. B) Effect of $K_{d,SK}$ modulation on the maximal basic cycle length (BCL) required for inducing voltage (solid lines) and $Ca^{2+}$ (dotted lines) alternans in the baseline cAF model. C) Effect of $K_{d,SK}$ modulation on the maximal BCL required for inducing delayed afterdepolarizations (DADs) in the baseline cAF model.

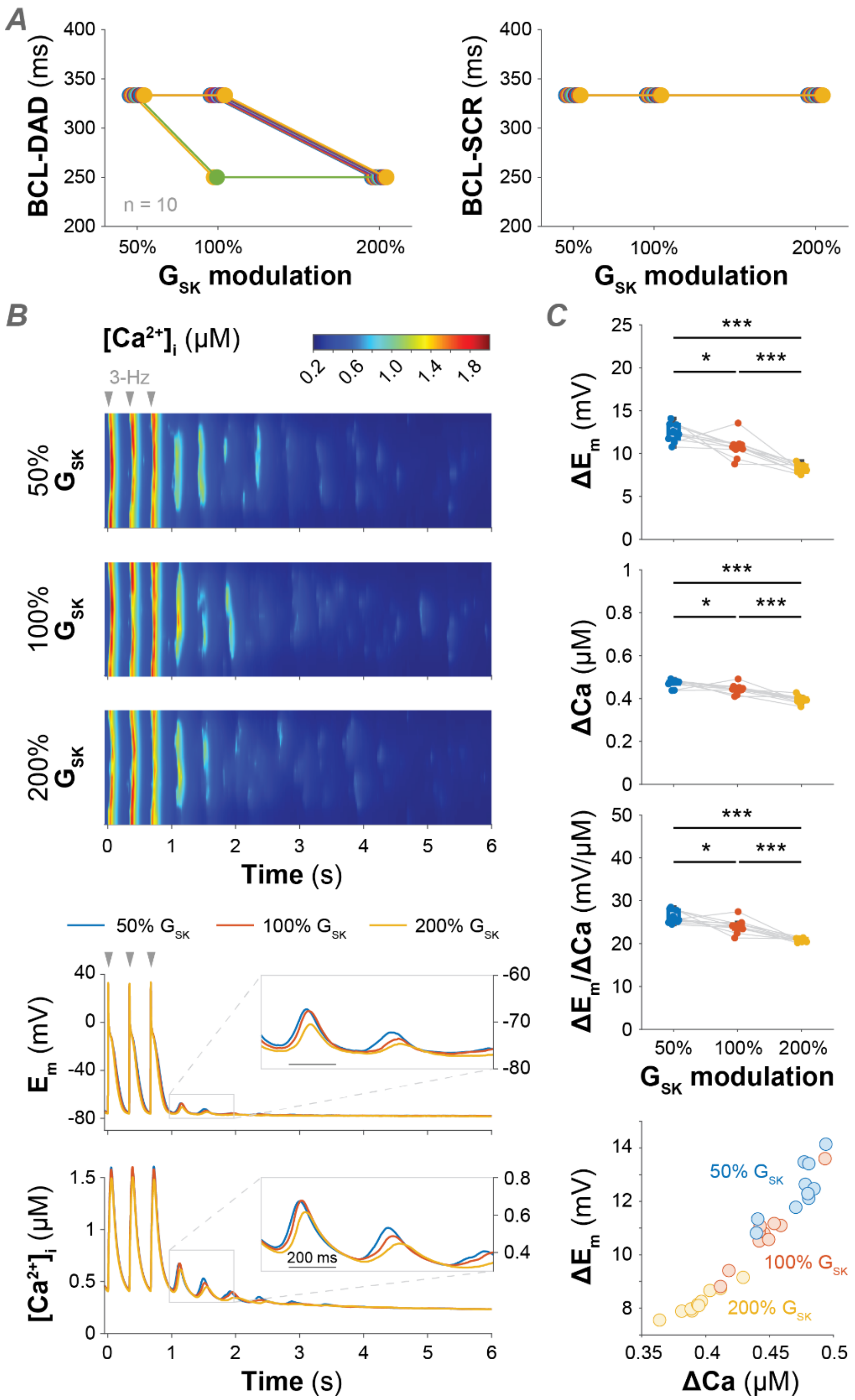


**Figure S4. Increasing SK channel conductance reduces the coupling between transmembrane potential and intracellular $Ca^{2+}$ also when SK affinity for intracellular $Ca^{2+}$ is increased.** We replicated the simulations shown in **Fig. 5** with increased SK channel affinity for intracellular $Ca^{2+}$ concentration (i.e., $K_{d,SK}$ reduced to 230 nM). A) Effect of changes in $G_{SK}$ on the maximal basic cycle length (BCL) inducing delayed depolarizations (DADs) and spontaneous $Ca^{2+}$ release events (SCRs) in the 3D atrial myocyte model. We used 10 mV and 300 nM as thresholds for DADs and SCRs, respectively. The simulations were repeated for 10 randomly generated tubular structures with sparse tubular densities. B) Top panels show representative transverse line scans of local cytosolic $Ca^{2+}$ concentration obtained pausing the electrical stimulation after a train of impulses at 3-Hz pacing for different $G_{SK}$ values. The last three paced beats are reported in the figure. Bottom panels report the time course of corresponding global cytosolic $Ca^{2+}$ concentration and membrane potential. C) Summary data resulting from simulating the protocol described in panel B in the 10 different tubular structures. Top panels show amplitude of $E_m$ and $[Ca^{2+}]_i$ oscillations ($\Delta E_m$ and $\Delta Ca$), and the $\Delta E_m/\Delta Ca$ ratio in function of different $G_{SK}$ values. Statistical analysis was performed by one-way ANOVA with Bonferroni correction (***: $p < 0.001$; **: $p < 0.01$; *: $p < 0.05$; ns: not significant). $\Delta E_m$ and $\Delta Ca$ are also compared against each other in the scatter plot reported in the bottom panel.

**Table S1. Definition of the Grandi *et al*. model parameters randomly perturbed to introduce variability in the population of models.**

| Parameter | Definition |
|---|---|
| $G_{Na}$ | Maximal conductance of the fast $Na^+$ current |
| $G_{NaL}$ | Maximal conductance of the late $Na^+$ current |
| $G_{NaB}$ | Maximal conductance of the background $Na^+$ current |
| $G_{to}$ | Maximal conductance of the transient outward $K^+$ current |
| $G_{Kr}$ | Maximal conductance of the rapidly activating $K^+$ current |
| $G_{Ks}$ | Maximal conductance of the slowly activating $K^+$ current |
| $G_{Kur}$ | Maximal conductance of the ultra-rapidly activating $K^+$ current |
| $G_{Kp}$ | Maximal conductance of the plateau $K^+$ current |
| $G_{K2P}$ | Maximal conductance of the two-pore-domain $K^+$ current |
| $G_{SK}$ | Maximal conductance of the small-conductance $Ca^{2+}$-activated $K^+$ current |
| $G_{K1}$ | Maximal conductance of the inward rectifier $K^+$ current |
| $G_{K,ACh}$ | Maximal conductance of the acetylcholine-sensitive $K^+$ current |
| $G_{CaL}$ | Maximal conductance of the L-type $Ca^{2+}$ current |
| $G_{CaB}$ | Maximal conductance of the background $Ca^{2+}$ current |
| $G_{ClCa}$ | Maximal conductance of the $Ca^{2+}$-dependent $Cl^-$ current |
| $G_{ClB}$ | Maximal conductance of the background $Cl^-$ current |
| $v_{NKA}$ | Maximal transport rate of the $Na^+/K^+$ ATPase |
| $v_{NCX}$ | Maximal transport rate of the $Na^+/Ca^{2+}$ exchanger |
| $v_{PMCA}$ | Maximal transport rate of the plasma membrane $Ca^{2+}$ ATPase |
| $v_{SERCA}$ | Maximal transport rate of the sarcoplasmic reticulum (SR) $Ca^{2+}$ ATPase |
| $v_{RyR,rel}$ | Maximal transport rate of the SR $Ca^{2+}$ release via ryanodine receptors (RyRs) |
| $v_{RyR,leak}$ | Maximal transport rate of the SR $Ca^{2+}$ leak via RyRs |
